# Social communication in mice – Are there optimal cage conditions?

**DOI:** 10.1101/011783

**Authors:** Allain-Thibeault Ferhat, Anne-Marie Le Sourd, Fabrice de Chaumont, Jean-Christophe Olivo-Marin, Thomas Bourgeron, Elodie Ey

## Abstract

Social communication is heavily affected in patients with neuropsychiatric disorders. Accordingly, mouse models designed to study the mechanisms leading to these disorders are tested for this phenotypic trait. Test conditions vary between different models, and the effect of these test conditions on the quantity and quality of social interactions and ultrasonic communication is unknown. The present study examines to which extent the habituation time to the test cage as well as the shape / size of the cage influence social communication in freely interacting mice. We tested 8 pairs of male mice in free dyadic social interactions, with two habituation times (20 min and 30 min) and three cage formats (rectangle, round, square). We tested the effect of these conditions on the different types of social contacts, approach-escape sequences, follow behavior, and the time each animal spent in the vision field of the other one, as well as on the emission of ultrasonic vocalizations and their contexts of emission. We provide for the first time an integrated analysis of the social interaction behavior and ultrasonic vocalizations. Surprisingly, we did not highlight any significant effect of habituation time and cage shape / size on the behavioral events examined. There was only a slight increase of social interactions with the longer habituation time in the round cage. Remarkably, we also showed that vocalizations were emitted during specific behavioral sequences especially during close contact or approach behaviors. The present study provides a protocol reliably eliciting social contacts and ultrasonic vocalizations in male mice. This protocol is therefore well adapted for standardized investigation of social interactions in mouse models of neuropsychiatric disorders.

## Introduction

Neuropsychiatric diseases affect heavily the social life of patients. In many cases, social communication is affected and becomes atypical. The patients get isolated in their social environment, due to a lack of interest for social interactions or atypical ways of interacting. Genetic studies of neuropsychiatric diseases led to the identification of several susceptibility genes for autism spectrum disorders (ASD) [1,2] or schizophrenia [3]. Mice remain one of the privileged mammalian animal models to study neuropsychiatric disorders, given the relative easiness of genetic modifications in this species.

Mice are social animals using olfactory, visual, tactile and acoustic signals to communicate [4–7]. Acoustic signals are one of the easiest communication signal types to measure and quantify, and they are considered as a reliable proxy to model social communication deficits [8,9]. Therefore, social interactions and vocal communication in mice are more and more often used as proxies to examine the validity of mouse models for neuropsychiatric diseases [10,11].

One issue in different tests used to evaluate social interactions and communication is to stimulate free social interactions without forcing them, i.e., favoring voluntary approaches instead of incidental contacts. The aim is to get a maximum of spontaneous affiliative social interactions. For this purpose, the shape of the test cage as well as the time the tested animal is habituated to it are supposed to be determinant (Figure 1A). A preliminary scanning of the literature of mouse models of ASD was not informative to highlight any relationship between cage size and social contact in free social interactions. Indeed, the duration of the recording had a major influence on the proportion of time spent in contact during that test. The shorter the test, the higher the percentage of time was spent in contact (Table S1).

**Figure 1:**
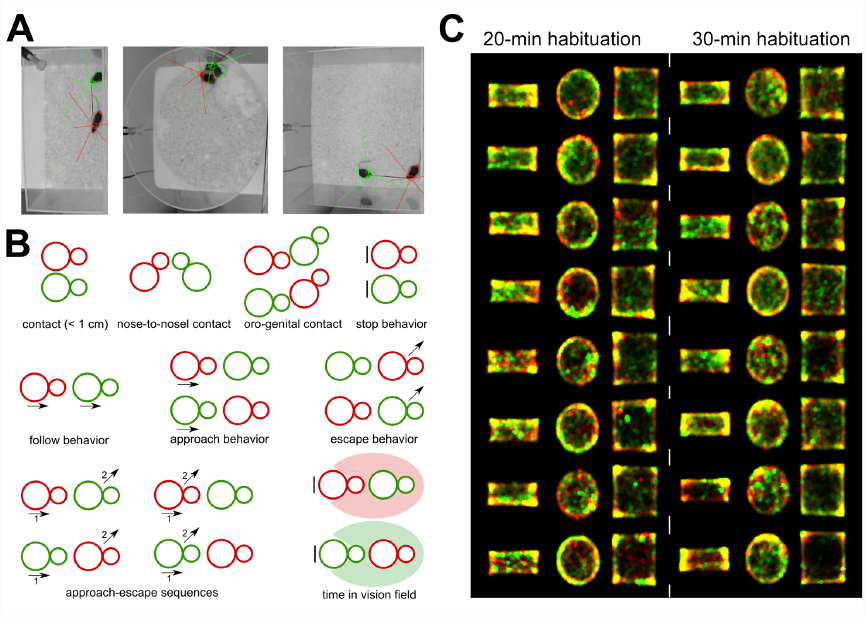
Methods used to examine the influence of cage shape/size on social interactions in mice. (A) Setting used to track with Mice Profiler the *occupant* male (red) and the *new-comer* male (green) during social interactions in a rectangle, a round and a square cage. (B) Behavioral events detected by Mice Profiler during social interactions between the *occupant* (red) and the *new-comer* (green). (C) Spatial occupation of the interaction cage by the *occupant* (red) and the *new-comer* (green) for the eight pairs of mice in the three cage types. Yellow traces represent the overlap of the two spatial occupations.

To avoid these influences, a standardized test was developed for the characterization of social deficits in mouse models of ASD and is now broadly used [11]. The three-chambered test has been designed in the laboratory of J. N. Crawley to evaluate the interest for a conspecific (constrained under a cup) as well as the preference for an unfamiliar conspecific over a familiar one [12]. Variables measured are usually the time spent in each compartment and the time spent sniffing the cups with the conspecifics (e.g., [13–16]. The advantage of this test is its standardization. Its major weakness is that social interactions are constrained, allowing only a very limited evaluation of social communication even for the sniffing time supposed to be the most representative of social interest [17]. This test allows the detection of social deficits; it gives however no qualitative indication about social interactions and represents only a quantitative estimation for social interest and conspecific recognition. A complementary test to counterbalance this weakness is the same-sex free dyadic interactions test [8,14,15,18,19]. Mice are left free to interact in a test cage. Both mice can be introduced together at the same time (situation not considered here), or one animal (the *occupant*, usually the tested mouse of a specific genotype) can be habituated to the test cage, while the second one is introduced later (the *new comer*, usually from a commercially available control strain). The last situation allows to distinguish between *occupant* and *new-comer*, i.e., the emitter and the receiver of most interactions, respectively. This test does not encompass as many standardized elements as the three-chambered test since animals are freely interacting and cannot be controlled. The analysis of this type of interactions is more time-consuming and more prone to subjective interpretations. New software nevertheless allow to reduce these subjective elements in the analyses (e.g., EthoVision XT from Noldus Information Technology, The Netherlands, or Mice Profiler from Icy software [20,21]). The elicited interactions are closer to ethological behaviors, even if the motivation of the animals is usually manipulated (through social isolation and habituation to the test cage for the *occupant*, and group housing for the *new comer*, situation considered in the present study). Data extracted allow a qualitative description of the interactions.

We designed the present study to test specifically the influence of the cage shape/size and the habituation time that vary in a controlled way. We aimed at testing whether same-sex social interactions occurring in a reasonably sized test cage with reduced habituation time are not forced in comparison with social interactions in larger cages with longer habituation time. We used the semi-automated module Mice Profiler from Icy software [20], that allows a comparison over a large panel of events, states and sequences of events within social interactions to get as much details as possible on these interactions in each condition.

We hypothesized that social interactions in the round cage will be the most different from the rectangle and the square cages. Indeed, their absence might lead to major modifications of the interactions. Given our previous experiments, two habituation times have been chosen close enough not to have a large corners in test cages appear to be very often used by animals (see Figure 1C), and therefore influence on social interactions. The reason of this comparison is very pragmatic: the shorter The reason of this comparison is very pragmatic: the shorter habituation time will be convenient to spare some time for each animal and to improve testing efficiency.

## Material and methods

Upon arrival, seven-week old C57BL/6J male mice (Charles River Laboratories, L’Arabesle, France) were housed in groups of four in a colony room maintained at 23±1°C on a 12:12 hour light/dark cycle, with lights on at 8:00 AM. Mice had access to food and water *ad libitum*.

One week after their arrival, the 48 males considered as *occupants* were socially isolated for 3 weeks before the experiments to increase their social motivation and reduce their aggressiveness [18,19]. The 12 mice used as *new-comers* were kept in groups of four males throughout the experiment. The experiments were conducted when animals were 11 weeks of age.

Behavioral experiments were approved by the ethical committee CETEA Institut Pasteur n°89. All experiments were conducted between 9:30 AM and 6:00 PM. Two pairs of mice were tested simultaneously with an opaque separation between the two cages. Pairs tested together underwent the same habituation time, but cage shape/size was balanced between right and left for each trial. Parameters tested in the experiment were the habituation time to the test cage (20 min or 30 min) and the shape and surface of the experimental Plexiglas cage (rectangle: 50 × 25 cm [1250 cm^2^], round: 50 cm diameter [1963.5 cm^2^], square: 50 cm side [2500 cm^2^]; height: 30 cm in all cases). The *occupant* mouse was left 20 min or 30 min for habituation in the experimental cage (100 lx; clean sawdust bedding [8]). After this time, an unfamiliar male mouse (the *new-comer*) was introduced. The two animals were allowed to freely interact for 8 min [8,18]. Social interactions were videotaped continuously (high-resolution CamTech Super-Hi-Res video camera; 25 fps). Ultrasonic vocalizations were recorded simultaneously (sampling frequency: 250 kHz, 16-bit accuracy). Audio recording hardware (UltraSoundGate 416-200, Condenser ultrasound microphone Polaroid/CMPA) and software (Avisoft SASLab Pro Recorder) were from Avisoft Bioacoustics (Berlin, Germany). Previous experiments suggested that ultrasonic vocalizations were mainly emitted by the *occupant*; the contribution of the *new-comer* (if any) was negligible [19]. At the end of the experiment, mice were returned to their respective home cages and none of the *occupants* were used again. Between each interaction test, the experimental cage was emptied from bedding, and new fresh bedding was used.

For ethical reasons, we tested and analyzed 8 pairs of mice per condition (habituation time: 20 min or 30 min; cage shape: rectangle, round, or square). Mice Profiler provides sufficiently detailed information to extract robust data with such a sample size for each condition (see supplementary information in [20]). The experimenters were blind of the conditions of the tested animals for data analyses (except for video scoring, where the experimenter could see the shape of the cage but was not aware of the habituation time). Social interactions were encoded on compressed video files (704 x 576 pixels; Figure 1A). Using the plugin Mice Profiler from Icy software [20], we measured several behavioral events (Figure 1B):

-time spent in contact (threshold: 1 cm),
-types of contacts: nose-to-nose, oro-genital from the *occupant*’s point of view and oro-genital from the *new-comer*’s point of view,
-follow behavior (i.e., the *occupant* follows the *new-comer*, both animals moving),
-approach behavior (i.e., the *occupant* approaches the *new-comer*, the *new-comer* approaches the *occupant*),
-escape behavior (i.e., the *occupant* escapes from the *new-comer*, the *new-comer* escapes from the *occupant*),
-approach-escape sequences (i.e., *occupant* approaches *new-comer* & *new-comer* escapes, *new-comer* approaches *occupant* and *occupant* escapes, *occupant* approaches *new-comer* & *occupant* escapes, *new-comer* approaches *occupant* & *new-comer* escapes),
-the time each animal spent in the vision field of the other one, both for the *occupant* and the *new-comer* when they are not moving.
-stop behavior (i.e., the *occupant* does not move, or the *new-comer* does not move).

We detected manually ultrasonic vocalizations with the software Avisoft SASLab Pro (Avisoft, Germany; FFT-length: 1024 points, 75 % overlap; time resolution: 0.853 ms; frequency resolution: 293 Hz; Hamming window). We measured the latency for the first ultrasonic vocalization as well as the total number of ultrasonic vocalization and their duration.

We used non-parametric statistical tests given the non-normal distribution of the data and the small sample-sizes (8 pairs of animals per condition). We used a Kruskal-Wallis rank sum test to evaluate the effect of the cage shape for each habituation time separately. When significant differences emerged, we used Wilcoxon-Mann-Whitney U-tests to identify where the difference stemmed from in 2-by-2 comparisons. To evaluate the effect of the habituation time for each cage shape separately, we used Wilcoxon-Mann-Whitney U-tests. All statistical results are compiled in the Table S2. All analyses were conducted with the computing and statistical software R(R Developmental Core Team 2009).

## Results

We tested the effect of the cage shape/size and the habituation time on different events and sequences of events occurring in affinitive social interactions between adult male mice. Results are presented for the first 4 min of the recordings. Results for the complete 8-min recordings are overall similar to the ones for the first 4-min recordings unless otherwise specified (data not shown; Table S2 for statistical analyses).

The habituation time to the test cage influenced the time spent in contact only in the round cage, with the time spent in contact being significantly longer after 30-min habituation than after 20-min habituation (Mann-Whitney U-test: U=10, p=0.021; Figure 2). There was no significant effect of the cage shape/size on the time spent in contact after 20-min habituation and after 30-min habituation (Figure 2).

**Figure 2:**
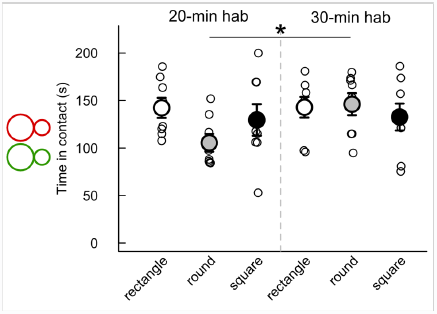
Limited influence of habituation time and cage shape/size on the time spent in contact. Time in contact (< 1 cm) was measured during the first 4 min of interaction after 20 min (left panel) or 30 min (right panel) habituation time in three cage types. (Wilcoxon-Mann-Whitney U-tests: n=8 pairs of mice per condition; ^*^: p<0.05; data are presented as mean +/- sem).

The effects of the habituation time and of the cage shape/size were also limited in the different types of contacts (Figure 3). There was a significant effect of habituation time in the round cage only on the total duration of nose-to-nose contacts (Mann-Whitney U-test: U=6, p=0.005; Figure 3A). This contact was significantly longer after 30-min habituation than after 20-min habituation in the round cage. There was no significant difference due to habituation time in the total duration of the *occupant* sniffing the ano-genital region of the *new-comer* and in the total duration of the *new-comer* sniffing the ano-genital region of the *occupant* (Figure 3B & C). Concerning cage shape/size, we did not find any significant effect neither in the 20-min habituation condition nor in the 30-min habituation condition on the total time spent in nose-to-nose contacts (Figure 3A), on the total time spent by the *occupant* sniffing the ano-genital region of the *new-comer* (Figure 3B), and on the total time spent by the *new-comer* sniffing the ano-genital region of the *occupant* (Figure 3C).

**Figure 3:**
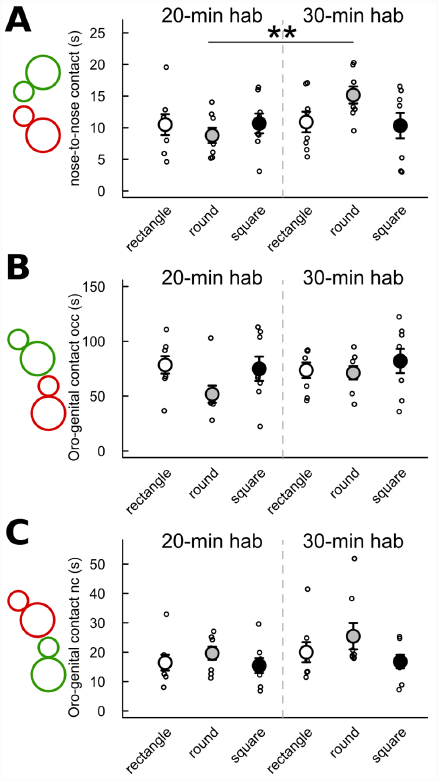
Limited effect of habituation time and cage shape/size on the duration of different types of contact. Contact (< 1 cm) was measured during the first 4 min of interaction after 20 min (left panel) or 30 min (right panel) habituation time in three cage types. (A) Time spent in mouth-to-mouth contact. (B) Time spent by the *occupant* (occ) sniffing the ano-genital region of the *new-comer*. (C) Time spent by the *new-comer* (nc) sniffing the ano-genital region of the *occupant*. (Wilcoxon-Mann-Whitney U-tests: n=8 pairs of mice per condition; ^*^: p<0.05, ^**^: p<0.01; data are presented as mean +/- sem).

The total duration of the *occupant* following the *new-comer* did not differ significantly according to the cage shape and the habituation time (all p-values >0.05; Figure 4A). No significant effect of habituation time and cage shape/size had been found on the duration of the stop behaviors (Figure 4B & C). No significant differences between the 20-min and the 30-min habituation conditions had been found in any cage shape/size for the number of occurrences of the *occupant* approaching the *new comer* (Figure 5A), the number of occurrences of the *new-comer* approaching the *occupant* (Figure 5B), and the number of occurrences of the *new-comer* escaping from the *occupant* (Figure 5D). Only the number of occurrences of the *occupant* escaping from the *new-comer* had been significantly higher and lower, respectively, after 30-min habituation than after 20-min habituation in the round cage (Mann-Whitney U-test: U=8 p=0.013) and in the square cage (Mann-Whitney U-test: U=55.5 p=0.016; Figure 5C). The effects of the cage shape/size were not significant in both the 20-min and the 30-min habituation conditions on the number of occurrences of the *occupant* approaching the *new-comer* (Figure 5A), the number of occurrences of the *new-comer* approaching the *occupant* (Figure 5B), and the number of occurrences of the *new-comer* escaping from the *occupant*

**Figure 4:**
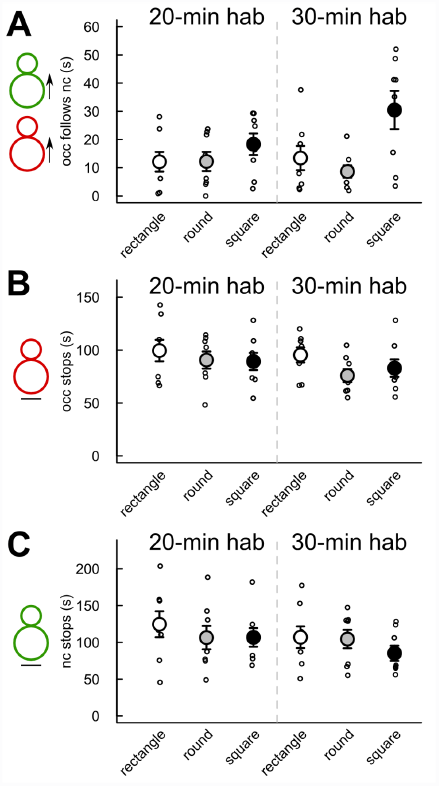
No significant effect of habituation time and cage shape/size on the duration of the follow and stop behaviors. Measures were taken after 20 min (left panel) or 30 min (right panel) habituation time in three cage types. (A) Time spent by the *occupant* (occ) following the *new-comer* (nc) in the first 4 min of interaction. (B) Time spent by the *occupant* (occ) not moving during the first 4 min of interaction. (C) Time spent by the *new-comer* (nc) not moving during the first 4 min of interaction. (Wilcoxon-Mann-Whitney U-tests: n=8 pairs of mice per condition; data are presented as mean +/- sem).

**Figure 5:**
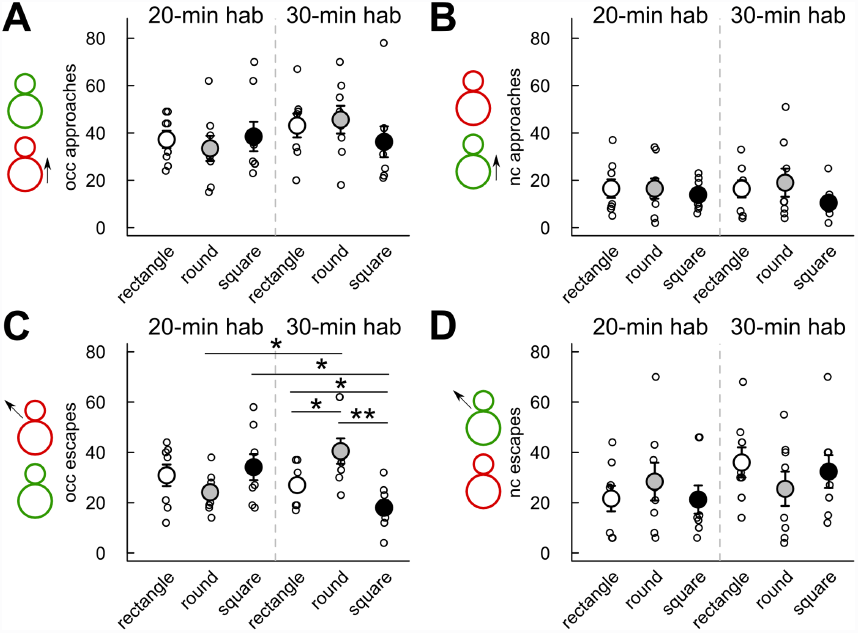
Limited effect of habituation time and cage shape/size on the number of approach or escape behaviors. Measures were taken during the first 4 min of interaction after 20 min (left panel) or 30 min (right panel) habituation time in three cage types. (A) Number of occurrences of the *occupant* (occ) approaching the *new-comer*. (B) Number of occurrences of the *new-comer* (nc) approaching the *occupant*. (C) Number of occurrences of the *occupant* (occ) escaping from the *new-comer*. (D) Number of occurrences of the *new-comer* (nc) escaping from the *occupant*. (Wilcoxon-Mann-Whitney U-tests: n=8 pairs of mice per condition; ^*^: p<0.05, ^**^: p<0.01; data are presented as mean +/- sem).

There had been only a significant effect of the cage shape/size in the 30-min habituation condition on the number of occurrences of the *occupant* escaping from the *new-comer* (Kruskal-Wallis test: Df=2, X=11.67, p=0.003; Figure 5D). Indeed, the *occupant* escaped from the *new-comer* significantly more frequently in the round cage in comparison with the rectangle cage (Mann-Whitney U-test: U=12, p=0.040) and in comparison with the square cage (Mann-Whitney U-test: U=60.5, p=0.003) after 30-min habituation. The *occupant* also escaped from the *new-comer* significantly more frequently in the rectangle cage in comparison with the square cage after 30-min habituation (Mann-Whitney U-test: U=51.5, p=0.045). Results for the duration of the same behavioral events were similar (data not shown).

Very limited influence of the cage shape/size and habituation time had been found on sequences of approach-escape behaviors (Figure 6). There had been no significant effect of the habituation time on the frequency of occurrence (Figure 6) and the duration (data not shown) of the three following types of sequences: “*occupant* approaches *new-comer* & *new-comer* escapes” (Figure 6A), “*new-comer* approaches *occupant* and *occupant* escapes” (Figure 6B), and “*new-comer* approaches *occupant* & *new-comer* escapes” (Figure 6D). Only a significant effect of habituation time had been found for the sequence “*occupant* approaches *new-comer* & *occupant* escapes” in the square cage (Figure 6C). Indeed, the sequence “*occupant* approaches *new-comer* & *occupant* escapes” occurred significantly less frequently in the 30-min habituation condition than in the 20-min habituation condition in the square cage (Mann-Whitney U-test: U=59.5, p=0.004). No significant differences in the frequency of occurrences (Figure 6) and in the duration (data not shown) of the four types of sequences emerged between the cage shapes in either the 20-min habituation condition. There had been nevertheless a significant effect of cage shape/size in the 30-min habituation condition for the sequence “*occupant* approaches *new-comer* & *occupant* escapes” (Kruskal-Wallis test: Df=2, X=6.33, p=0.042), which occurred significantly more frequently in the round cage in comparison with the square cage (Mann-Whitney U-test: U=54, p=0.023; Figure 6C).

**Figure 6:**
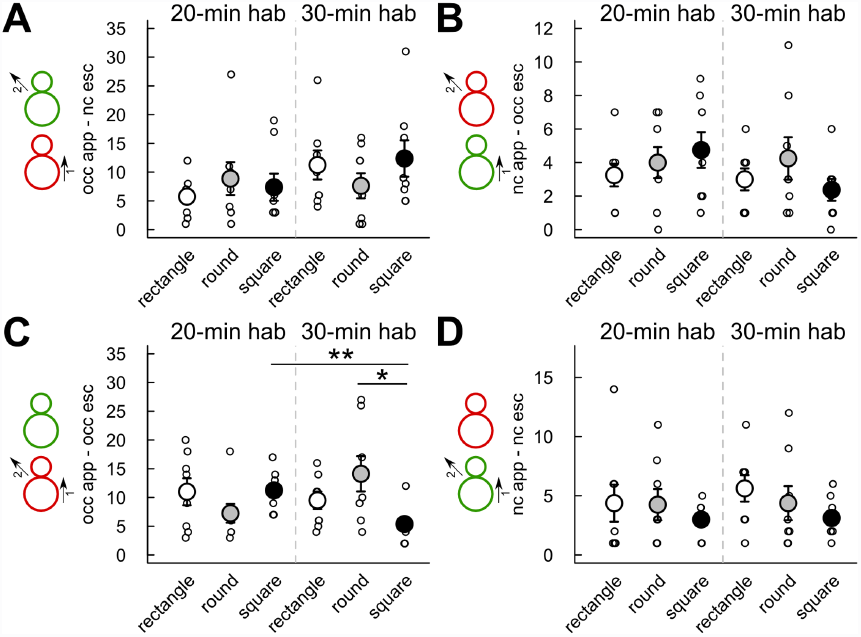
Limited effect of habituation time and cage shape/size on the number of occurrences of different types of approach-escape sequences. Measures were taken during the first 4 min of interaction after 20 min (left panel) or 30 min (right panel) habituation time in three cage types. (A) Number of occurrences of the sequence “*occupant* approaches *new-comer* & *new-comer* escapes”. (B) Number of occurrences of the sequence “*new-comer* approaches *occupant* and *occupant* escapes”. (C) Number of occurrences of the sequence “*occupant* approaches *new-comer* & *occupant* escapes”. (D) Number of occurrences of the sequence “*new-comer* approaches *occupant* & *new-comer* escapes”. (Wilcoxon-Mann-Whitney U-tests: n=8 pairs of mice per condition; ^*^: p<0.05, ^**^: p<0.01; data are presented as mean +/- sem).

We also examined the effect of cage shape/size and habituation time on the time spent by the animals in the vision field of each other when the animals are not moving. The *occupant* kept the *new-comer* in its vision field significantly longer after 20-min habituation than after 30-min habituation in the round cage only (Mann-Whitney U-test: U=51, p=0.049; Figure 7A). No significant effect of habituation time had been found on the time the *new-comer* kept the *occupant* in its vision field (Figure 7B). The cage shape/size did not appear to have a significant influence on the time spent by each animal in the vision field of the other one during stop situations for both the 20-min habituation condition and the 30-min habituation condition during the first 4 min of interactions (Figure 7). In contrast, over the 8 min of interactions, there was a significant effect of cage shape/size on both the time the *occupant* kept the *new-comer* in its vision field (Kruskal-Wallis test: Df=2, X=8.14, p=0.017) and the time the *new-comer* kept the *occupant* in its vision field (Kruskal-Wallis test: Df=2, X=7.27, p=0.026). Both measures were more elevated in the round cage in comparison with the rectangle cage (Mann-Whitney U-tests: U=5, p=0.003; U=11, p=0.028; data not shown).

**Figure 7:**
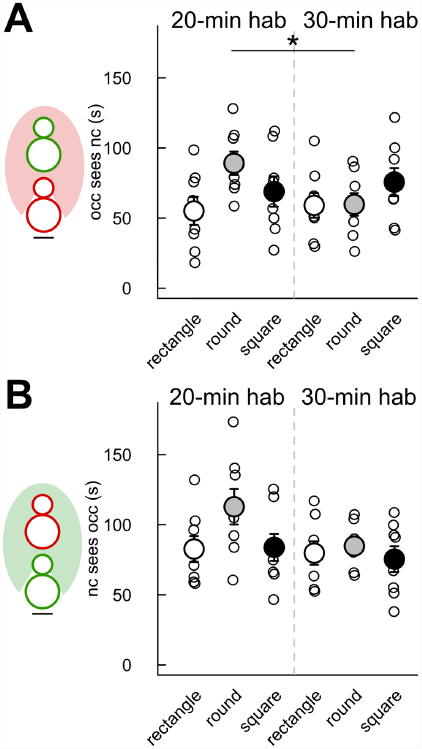
Limited effect of habituation time and cage shape/size on the time spent by one individual in the vision field of the other one. Measures were taken during the first 4 min of interaction after 20 min (left panel) or 30 min (right panel) habituation time in three cage types. (A) Time spent by the *new-comer* in the vision field of the *occupant*. (B) Time spent by the *occupant* in the vision field of the *new-comer*. (Wilcoxon-Mann-Whitney U-tests: n=8 pairs of mice per condition; ^*^: p<0.05; data are presented as mean +/- sem).

Finally, we tested the effect of the cage shape/size and the habituation time on the latency for the first ultrasonic vocalization, the number of ultrasonic vocalizations and the mean duration of the calls. No significant differences emerged between the habituation time conditions for each cage shape and between cage shapes for each habituation time condition for the latency of the first ultrasonic vocalization (Figure 8A), the number of ultrasonic vocalizations emitted per minute (Figure 8B) and the mean duration of ultrasonic vocalizations (Figure 8C).

**Figure 8:**
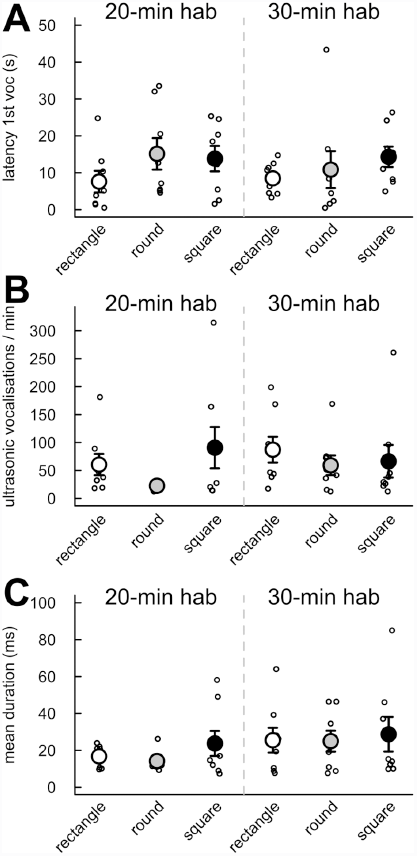
No significant effect of habituation time and cage shape/size on the vocal behavior of the pairs of interacting mice. PMeasures were taken during the first 4 min of interaction after 20 min (left panel) or 30 min (right panel) habituation time in three cage types. (A) Latency for the first ultrasonic vocalization. (B) Number of ultrasonic vocalizations emitted by minute. (C) Mean duration of ultrasonic vocalizations. (Wilcoxon-Mann-Whitney U-tests: n=8 pairs of mice per condition; data are presented as mean +/- sem).

The behavioral events during which ultrasonic vocalizations were recorded did not vary significantly with habituation time or cage shape/size (Figure 9). In all conditions, between 30 % and 50 % of ultrasonic vocalizations were emitted during social contacts. Many vocalizations were recorded when the *occupant* was behind the *new-comer*, and much fewer in the reverse situation. This would explain why very few vocalizations were recorded during nose-to-nose contacts and when the *new-comer* was sniffing the ano-genital region of the *occupant*, while many vocalizations were recorded when the *occupant* was sniffing the ano-genital region of the *new-comer*. For vocalizations not emitted during contact, a high proportion of vocalizations recorded when the *occupant* was approaching the *new-comer* (with either the *new-comer* or the *occupant* escaping). In contrast, much fewer vocalizations were recorded when the *new-comer* was approaching the *occupant* (with either the *occupant* escaping, or the *new-comer* escaping).

**Figure 9:**
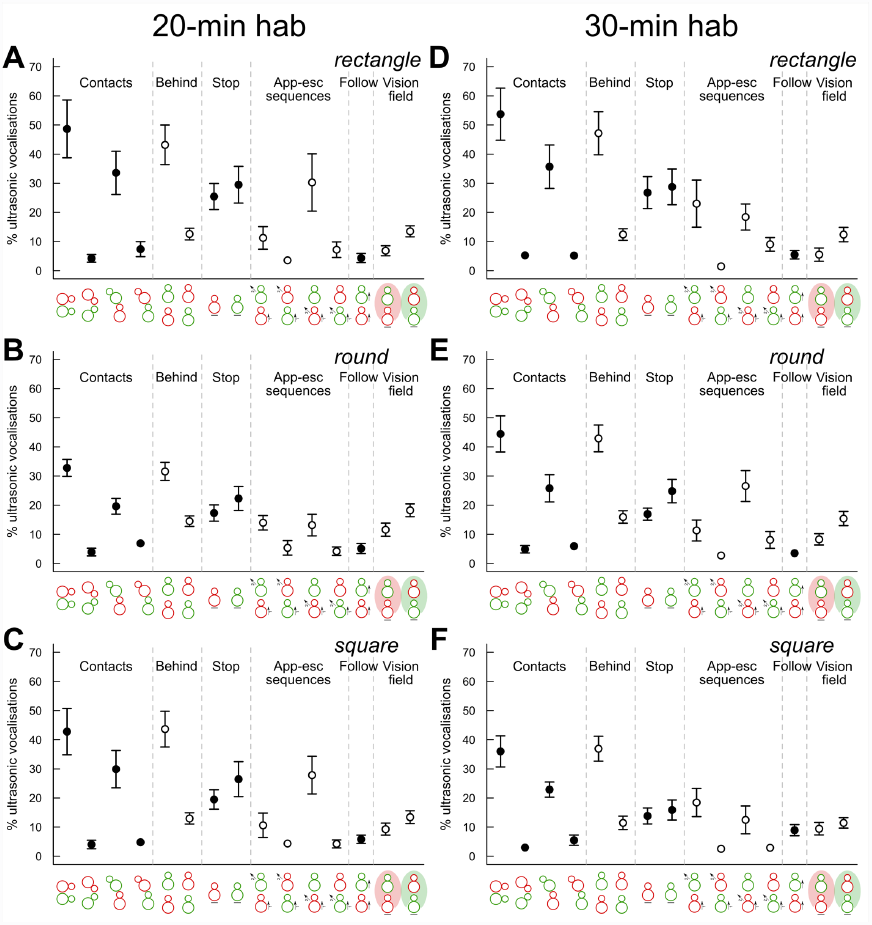
Contexts of ultrasonic vocalization emission during social interactions. Proportion of ultrasonic vocalizations emitted during social events during the first 4 min of interactions (A) after 20 min habituation in the rectangular cage, (B) after 20 min habituation in the round cage, (C) after 20 min habituation in the square cage, (D) after 30 min habituation in the rectangular cage, (E) after 30 min habituation in the round cage, (F) after 30 min habituation in the square cage. Social events are presented in the following order: contact < 1 cm, nose-to-nose contact, *occupant* sniffing ano-genital region of *new-comer*, *occupant* sniffing ano-genital region of *new-comer*, *occupant* behind *new-comer*, *new-comer* behind *occupant*, *occupant* not moving, *new-comer* not moving, *occupant* approaches *new-comer* & *new-comer* escapes, *new-comer* approaches *occupant* and *occupant* escapes, *occupant* approaches *new-comer* & *occupant* escapes, *new-comer* approaches *occupant* & *new-comer* escapes, *occupant* following *new-comer*, *new-comer* in the vision field of *occupant*, *occupant* in the vision field of *new-comer* (n=8 pairs of mice per condition; data are presented as mean +/- sem).

## Discussion

Using different settings for cage shape/size and habituation time, we ascertained mouse social behaviors and sequences of social behaviors, as well as ultrasonic vocalizations. We found very little difference caused by the cage shape/size in these features (Figure 10). Only the habituation time had some minor effect on social behaviors most often in the round cage, with the longer habituation time favoring social interactions.

**Figure 10:**
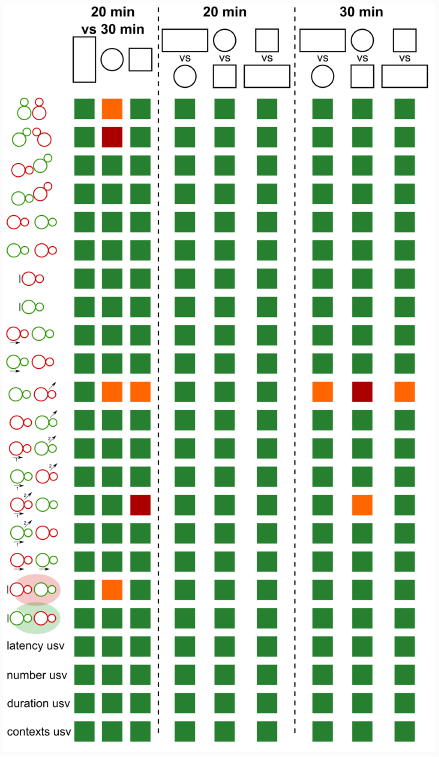
Limited effects of habituation time and cage shape/size on behavioral events during male mouse social interactions. Most comparisons were non-significant (green boxes: p > 0.05), while only a few effects of habituation time and cage shape/size were detected (orange boxes: p < 0.05, red boxes: p < 0.01).

Previous studies only analyzed the influence of housing conditions on the behavior of the animals (e.g., individually ventilated cages vs. filter-top cages [22], transparent walls vs. cages without view on the room and other cages [23]). In our study, we examined the direct influence of the cage shape/size during a social interaction test. We did not identify major effects of the cage shape/size and the habituation time on the several variables describing social interactions. The present study therefore suggests that the use of a 50 x 25 cm rectangular cage with a 20 min habituation time does not force social interactions, i.e., favor voluntary social approaches. This forcing of social interactions would come from a too small space. In such conditions, mice would come in contact incidentally much more often than in larger spaces during the exploration of the cage. What is still unknown and would need further investigation is to what extent interactions would be forced in a smaller space. Further study should also compare the quantity and quality of spontaneous social interactions with and without habituation to the test cage for the *occupant*.

We would have expected major differences in the round cage in comparison with the square and the rectangle ones given the absence of corners. Indeed corners might represent stop zones, where animals can get stuck. We thought this would lead to major differences for instance in approach/escape sequences. Such major differences were not observed. Only a slight increase of escape behaviors of the *occupant* from the *new-comer* has been found in the round cage in comparison with other cage types. This suggests that corners in test cages are not leading to major perturbations of ethological social sequences in mice.

The Mice Profiler module of Icy software from de Chaumont and colleagues [20] was originally tested on mice interacting with a similar protocol as ours, with a rectangular cage and 30 min habituation to the test cage. We compared our data within the rectangular cage and after 30 min habituation to these data for C57Bl/6J male mice. The duration of the follow behavior was slightly shorter in our experiment in comparison with the de Chaumont and colleagues’ study both during the first 4 min of interaction and during the 8 min of interaction [20]. The duration of nose-to-nose contact was lower in our study in comparison with the de Chaumont study, while the total duration of oro-genital sniffing (from both the *occupant* and the *new-comer*) was similar between the two studies during the first 4 min of interaction. Overall, most discrepancies were found for the nose-to-nose contact, while data for other behaviors were similar. The fact that mice from the present study were isolated for 3 weeks and those from the de Chaumont study for 4 weeks might explain some of the differences between the durations of the follow behavior and of the types of contacts.

Ultrasonic vocalizations were recorded in all conditions, confirming that two interacting males vocalize together and do not show overt aggressive behaviors during 4 min (and even 8 min) of interactions if the *occupant* has been isolated for 3 weeks before the test. Moreover, a similar amount of ultrasonic vocalizations was recorded in the three cage types. This absence of difference in the amount of vocalizations detected suggests that even in the largest cages (square and round cages), where the microphone has to cover a large surface, the level of detection of ultrasonic vocalizations is not different from the smallest cage (rectangle cage). This indication of the recording surface of the ultrasonic microphone used will be helpful to design new recording cages for 24 h continuous recordings of the animals in complex environments, including more than two individuals.

For the first time, we manage to correlate the emission of ultrasonic vocalizations and the detailed social events extracted from Mice Profiler. We confirmed that ultrasonic vocalizations are not emitted randomly and therefore provide further support for their role in social communication. At the moment, it is still not possible to identify clearly the emitter. The fact that ultrasonic vocalizations are rarely emitted during nose-to-nose contact could justify using triangulation to localize the emitter since the noses of the two animals are usually not close to one another during ultrasonic vocalization emission.

Ultrasonic vocalizations are supposedly mostly emitted by the *occupant* during social contacts (especially ano-genital sniffing) and during approach behaviors. The present study also suggested an effect of the emission of ultrasonic vocalizations on the outcome of an approach. The escape behavior of the *new-comer* or the *occupant* was often correlated with the emission of ultrasonic vocalizations. When the *occupant* approaches the *new-comer*, the later is often less likely to escape and end the contact if more vocalizations were emitted. New social experiments with long term continuous recordings of spontaneous interactions and playback studies should allow to shed more light on the functions of these ultrasonic vocalizations, and more specifically to detect any sequences triggering specific behaviors in the receiver.

Overall, the present study suggest that a rectangular cage (50 x 25 cm) with 20-min habituation to the test cage is sufficient to elicit non-aggressive interactions between an isolated “*occupant*” adult male and a socially-housed “*new-comer*” adult male. A large amount of ultrasonic vocalizations can even be recorded in these conditions. Such a protocol will be useful in a better evaluation of social communication deficits in mouse models of neuropsychiatric disorders.

## Acknowledgments

We thank Guillaume Dumas for valuable comments on the manuscript.

## Financing

This work was supported by the Fondation de France; by the ANR FLEXNEURIM [ANR09BLAN034003]; by the ANR [ANR- 08-MNPS-037-01–SynGen]; by Neuron-ERANET (EUHF-AUTISM); by the Fondation Orange; by the Fondation FondaMentale; by the Fondation Bettencourt–Schueller. The research leading to these results has also received support from the Innovative Medicines Initiative Joint Undertaking under grant agreement no. 115300, resources of which are composed of financial contribution from the European Union’s Seventh Framework Program (FP7/2007-2013) and EFPIA companies’ in kind contribution.

## Authors’ contribution

EE and TB conceived and designed the experiments. EE and AMLS collected the data. ATF, AMLS, FDC, JCOM, TB and EE analyzed the data and wrote the paper.

